# Reinforcement learning when your life depends on it: a neuro-economic theory of learning

**DOI:** 10.1101/2024.05.08.593165

**Authors:** Jiamu Jiang, Emilie Foyard, Mark C.W. van Rossum

**Affiliations:** University of Nottingham, Nottingham, UK

**Keywords:** reinforcement learning, learning and memory, metabolism, insects, computational modeling

## Abstract

Synaptic plasticity enables animals to adapt to their environment, but memory formation can consume a substantial amount of metabolic energy, potentially impairing survival. Hence, a neuro-economic dilemma arises whether learning is a profitable investment or not, and the brain must therefore judiciously regulate learning. Indeed, in experiments it was observed that during starvation, Drosophila suppress formation of energy-intensive aversive memories. Here we include energy considerations in a reinforcement learning framework. Simulated flies learned to avoid noxious stimuli through synaptic plasticity in either the energy expensive long-term memory (LTM) pathway, or the decaying anesthesia-resistant memory (ARM) pathway. The objective of the flies is to maximize their lifespan, which is calculated with a hazard function. We find that strategies that switch between the LTM and ARM pathways based on energy reserve and reward prediction error, prolong lifespan. Our study highlights the significance of energy-regulation of memory pathways and dopaminergic control for adaptive learning and survival. It might also benefit engineering applications of reinforcement learning under resources constraints.

---

Learning allows animals to adapt to their surroundings, evade dangers, and enhance survival prospects. However, learning itself comes at a cost as it requires considerable amounts of metabolic energy. For instance, experiments have shown that fruit flies that learn a classical conditioning task perish 20% faster when subsequently starved compared to starved control flies (Mery and Kawecki, 2005). When they are not starved, flies strongly increase their food intake after learning (Plaçais and Preat, 2013).

In Drosophila memory is expressed in (at least) two distinct pathways, that are believed to be mutually exclusive (Isabel et al., 2004). The Long Term Memory (LTM) pathway requires a lot of energy but yields persistent memory. Conversely, the Anesthesia Resistant Memory (ARM) pathway is thought to require negligible amounts of energy, as its expression does not significantly affect lifetime (Mery and Kawecki, 2005). However, ARM memory typically dissipates within four days (Tully et al., 1994). Notably, in aversive conditioning protocols flies halt energy-demanding LTM formation when starved (Plaçais and Preat, 2013).

As learning comes at a cost, a neuro-economic dilemma arises whether learning is a profitable investment or not. Yet, the energy requirements of learning have thus far been mostly overlooked in the computational community. The situation can be compared to the human dilemma whether or not to spend money on education: typically investment in education will pay off financially, but only if the life expectancy is long enough and bankruptcy can be avoided.

Here we examine the energy cost-benefit of learning on expected survival, and compare learning strategies that maximize survival during an aversive conditioning protocol. We introduce a hazard framework to examine the trade-off between the energy expenditure required learning and encountering hazardous stimuli. Learning to evade aversive stimuli decreases the stimulus hazard, but the energy expenditure associated with learning increases the starvation hazard. The objective for the flies is to maximize their lifetime by employing either the LTM or the ARM memory pathways. We propose a strategy that switches between ARM and LTM pathways depending on the current energy reserve and the reward prediction error. This strategy robustly increase life-time across a number of stimulus protocols.

## Model: Hazard framework

Most biological reinforcement learning studies assume that animals seek to maximize total reward and minimize punishment. The tacit assumption is that this improves biological fitness. It is then common to compare behavior to reward maximizing policies (e.g. Beron et al., 2022), often without regards for metabolic cost of implementing and updating the policy. Here, however, we directly assume that the optimal policy maximizes survival, i.e. the life time of the organism. Because learning requires energy, the policy needs to balance avoidance of a hazardous stimulus against the expenditure of energy by learning.

To examine this trade-off we use a hazard function approach. Hazard functions were originally developed in life insurance to calculate the probability that policy holders would die; they are also used in failure analysis and healthcare (e.g. Modarres et al., 1999; Clark et al., 2003). In computational neuroscience hazard functions have been used to model the probability that a neuron fires a spike (Gerstner et al., 2014). Despite being a natural approach, a hazard framework has to our knowledge not been used before for reinforcement learning problems.

Using a discrete time formulation, the hazard function *h*(*t*) (0 ≤ *h*(*t*) ≤ 1) specifies at any time the probability to die within a time unit. The probability to have a life time *t* is given by the probability of surviving all previous time-steps and perishing at time *t*. For a constant hazard *h*(*t*) = *h*, one finds that *P* (*t*) = (1 − *h*)^*t−*1^*h*, Fig. 1a. The life time distribution is in this case exponential with mean lifetime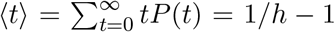. For time-varying hazards, the probability to have a life-time *t* is *P* (*t*) = *S*(*t*) − *S*(*t* + 1), where 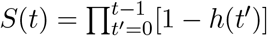 is the survival function to survive until time *t*. The mean life time follows as

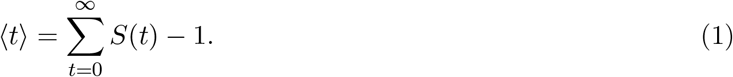

**Figure 1.**
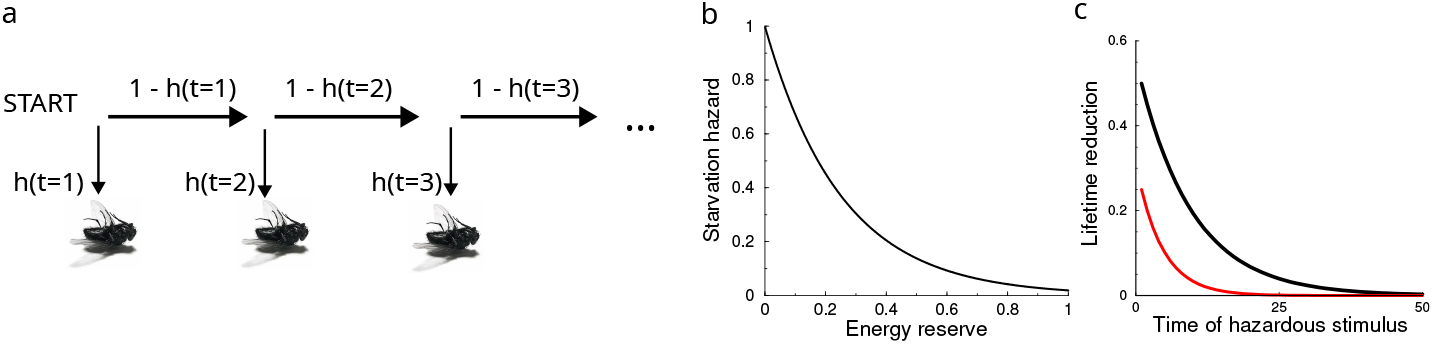
Hazard framework a) Illustration of hazard formulation. At each day the fly has a probability *h*(*t*) to die, or to survive to the next day. The hazard is determined by the fly’s metabolic energy reserve and the stimuli it encounters. The hazard has two components: starvation hazard and hazard from approaching the noxious stimulus. b) Assumed relation between the normalized energy reserve of the fly and its starvation hazard. The hazard increases exponentially at low energy. Note that even at maximal energy, there is a background hazard. c) Hazard framework leads to discounting. The reduction in lifetime due to an additional hazard versus the time of this extra hazard. Future hazards are exponentially less important than immediate ones. When the expected lifetime is shorter (red curve), the discounting is stronger, i.e. decay is faster. Baseline hazard: 0.1 (black), and 0.2 (red); hazard of stimulus in both cases 0.05.

In the following we measure time in days, and so the hazards have units ‘per day’.

The total hazard can include factors such as the internal state of the animal, as well as external stimuli and environmental factors. We consider two hazards: First is the hazard from starvation, which increases when the metabolic energy reserve *M* (*t*) diminishes. We assume that the energy reserve *M* (*t*) is positive and saturates at 1, corresponding to about 10 Joule (Girard et al., 2023). Although it would be straightforward to determine dependence of hazard on energy reserve experimentally, we are not aware of such experiments. Therefore we assume a steep increase at low energy levels, Fig. 1b,

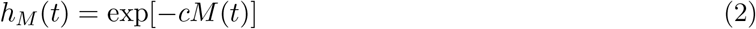

We calibrate *c* by using that well-fed flies (*M* = 1) have a lifespan of some 50 days (Linford et al., 2013), i.e. *c* = 3.9. Note that the hazard formulation includes the non-stochastic case where flies die if and only if the energy reaches zero. Hereto one would set the hazard *h*_*M*_ (*t*) to zero whenever *M >* 0, and to one otherwise.

Second, there is a hazard associated to approaching the aversive stimulus. Although laboratory experiments generally involve non-lethal shock stimuli, in a natural environment such shocks could potentially forebode a life threatening event, for instance the presence of a predator. We denote this hazard *h*_*s*_.

Being probabilities, hazards from different sources add up as *h*_Σ_(*t*) = 1 − [1 − *h*_*s*_(*t*)][1 − *h*_*M*_ (*t*)]. (In the limit of small *h*_*i*_ or, equivalently, the continuum limit, this reduces to a regular sum.)

Interestingly, the hazard framework automatically leads to reward discounting – a core feature added by hand to many RL models to express that immediate rewards are preferable to future rewards. In the hazard formulation rewards and hazards that are far in the future will hardly impact the lifetime. Instead it is important to minimize hazards early on. To illustrate discounting in a simple scenario, assume a constant permanent hazard and that at a certain time an additional hazard is introduced, active during one time-step only. The lifetime is reduced most if the hazard occurs immediately, whereas stimuli far in the future have no effect on the lifetime, Fig. 1b. For a constant background hazard, the discounting can be shown to be exponential. Furthermore, when the energy reserve is low and the expected lifetime shorter, the discounting is stronger, Fig. 1b (red curve).

Hazard typically also increases with age, however we assume that the experiments are so drastic that age dependence of the hazard can be ignored (“biologically immortality”) or averages out. In more detailed models such effects could be included. For instance, such models should find that expensive LTM learning is less beneficial for aged animals with little expected life-time left.

## Model: Network Design

We implemented a network reflecting the Drosophila brain’s anatomical structure, and a complementary feedback network associated with reinforcement, Fig. 2. In Drosophila aversive conditioning experiments, an odor (conditioned stimulus, CS) is paired with a shock (unconditioned stimulus, US). By repeating exposure to the CS-US pairs a few times, the flies learn to avoid the odor, as can be subsequently tested in a T-maze. The underlying circuitry, involving sensory encoding Kenyon Cells (KCs) and action-driving Mushroom Body Output Neurons (MBONs), is relatively well understood (Tempel et al., 1983; Tully et al., 1994; Bennett et al., 2021).

**Figure 2.**
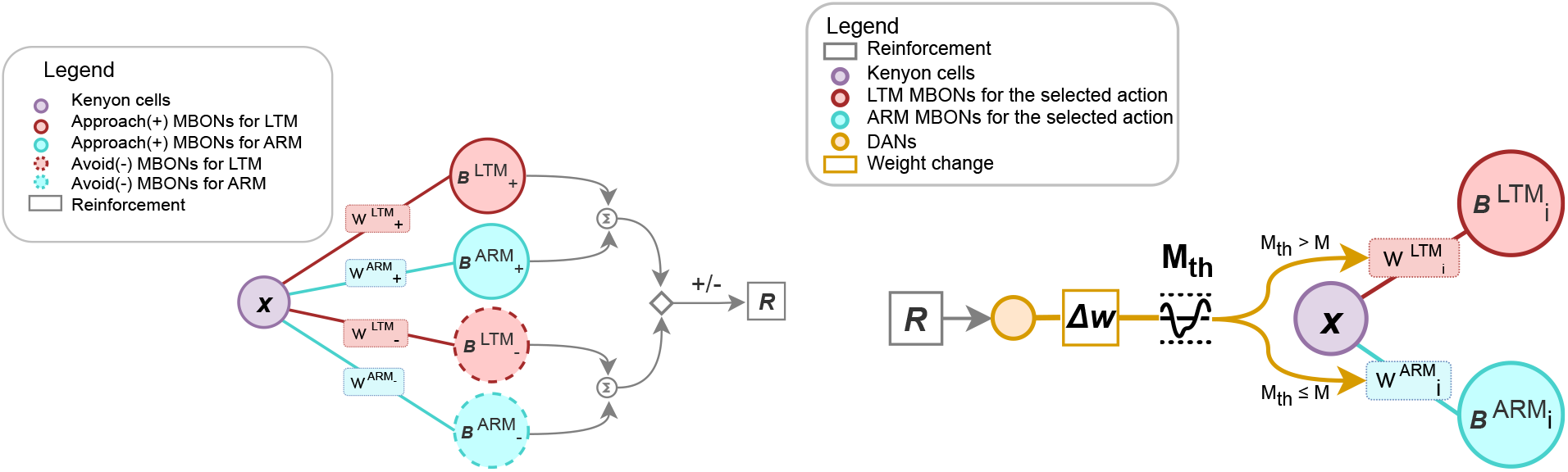
Schematic of the learning network rooted in the Drosophila brain anatomy. The left panel demonstrates the feed-forward Decision-making network, complemented by the right panel showing the feedback Energy-Adaptive learning mechanism, steered by reinforcement signals. The weight change, denoted by Δ*w*, is modulated by the reinforcement. The subscript represents the action of the current trial, either approach (+) or avoid (−).

The network comprises a population of sensory KCs that represent the odor signal, which subsequently drives the Mushroom Body Output Neurons (MBONs) that determine behavior, Fig. 2 left. The firing rate of the KC population is denoted *x*.

The activities of the MBONs are split up in the ARM and LTM pathways (see below). Each pathway is modeled as a linear neuron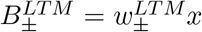, and 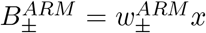, where ± indicates approach (+) and avoidance (−) behaviors, and the parameters 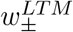 and 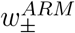 denote the synaptic strengths from the KCs to the MBONs. Given the additive nature of MBON signals (Aso et al., 2014b), we posit that total neuronal activity driving the approach and avoidance behaviors result from the sum of the ARM and LTM components. Hence

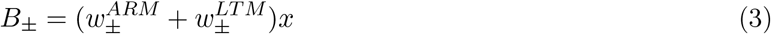

The total weight for approach and avoidance behaviors is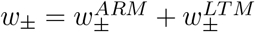.

Winner-Take-All competition between the two MBON neuron populations determines the fly’s action. The competition process is not explicitly modeled, but could reflect lateral inhibition and attractor dynamics. We assume that the neural processing and resulting decision making is noisy. (Otherwise, even the smallest imbalance would fully determine the decision). This randomness also means that the organism does not fully commit to avoiding even the smallest hazard, but keeps exploring as well. Assuming independent Poisson spike-time variability, the input to the decision making neurons has a variance equal to the mean input. At sufficient high rates this is well approximated by normal distribution with a variance equal to the mean. The probability to avoid *P*_*−*_ is a sigmoidal function of the difference in activities *B*_+_ and *B*_*−*_

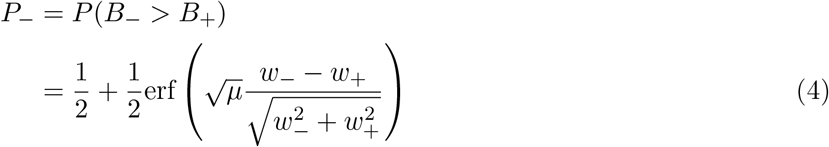

The mean *µ* of *x* can be extracted from the observation that when learning is saturated the performance corresponds to about *P*_*−*_ = 0.925 (Tully et al., 1994). Using *w*_*−*_ = 1 and *w*_+_ = 1*/*2 (see below), this yields *µ* = 10.3. The *µ* is the average number of spikes the MBON neuron receives from the sensory neurons within one integration period (e.g. 103Hz in 100ms); encouragingly it is similar to the value used in Bennett et al. (2021).

### Reward driven plasticity

The reward when approaching (+) the aversive stimulus is negative and denoted *R*_+_, without loss of generality we set *h*_*s*_ = −*R*_+_. That is, the punishment is expressed as its hazard. The reward for avoiding the stimulus, *R*_*−*_, is set to 0. In the MB of Drosophila, reinforcement-related signals are encoded by dopamine neurons (DANs) (Aso et al., 2014a), and these DAN signals modulate the plasticity of the synapse connecting KCs to MBONs (Cohn et al., 2015; Bennett et al., 2021). The synaptic strength associated with the selected behavior is updated based on the discrepancy between the reward from the current trial *R*_*±*_(*t*) and the expected reward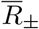, also known as the reward prediction error. The synaptic weight modification is

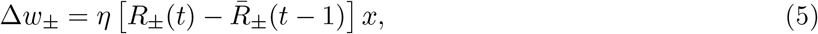

where *η* is the learning rate. In line with experiments (Hige et al., 2015), the learning according to Eq.5, occurs through depression of the approach action, rather than a strengthening of the avoid action. The learning rate was calibrated by using that in Tully et al. (1994) after a single cycle of learning, avoidance performance was *P*_*−*_ = 0.85, which corresponds to *w*_*−*_ = 0.8. Using *h*_*s*_ = 0.1 we find *η* = 0.6. As the performance early after learning through LTM and ARM are similar, the same learning rate was used for both ARM and LTM learning.

The 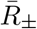 in Eq.5 is the running average of the reward of either action. The expected rewards are initialized at zero at the beginning of the simulation. The expected reward is updated when that action is chosen, otherwise it decays to zero

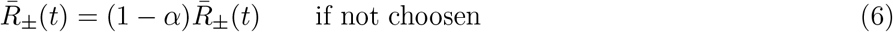

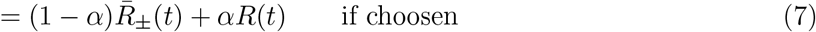

where *α* = 1 − *e*^*−*1*/τR*^, and the decay time constant of the average, *τ*_*R*_, is set equal to the ARM decay (below).

### Synaptic plasticity pathways

Experiments show that ARM and LTM memory formation are mutual exclusive (Isabel et al., 2004). Hence the synaptic weight changes given by Eq. 5 are expressed in either LTM or ARM weights. Updating the weight in the ARM pathway 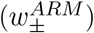 comes at negligible metabolic cost (Mery and Kawecki, 2005; Plaçais and Preat, 2013). However, the ARM weights decay over time, so that the update equation reads

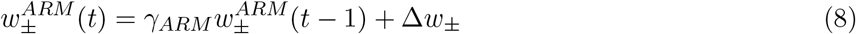

Here *γ*_*ARM*_ is the ARM decay rate. To estimate its value, we use that the data in Tully et al. (1994), where flies where exposed to massed training and the memory decay was measured. In four days the probability for the correct action decayed from *P*_*−*_ = 0.925 to *P*_*−*_ = 0.525 (in terms of the performance index used there, from 85% to 5%). In the model the memory extinction is found by substitution of Eq. 8 in Eq. 4. A fit yields *γ*_*ARM*_ = 0.34.

When, in contrast, LTM is expressed, the weight updates do not decay

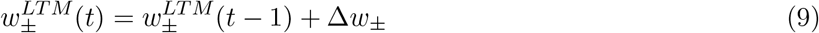

However, LTM is metabolically costly (Mery and Kawecki, 2005). We examine two abstract energy models. The first assumes that the metabolic energy cost of LTM formation decreases the energy reserve by an amount proportional to the weight change (Li and van Rossum, 2020)

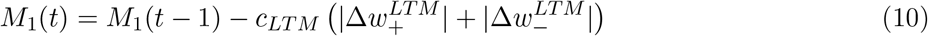

The parameter *c*_*LT M*_ denotes the energy cost of LTM. In experiments LTM before starvation reduced the survival time in female flies from 26 to 22 hrs (Mery and Kawecki, 2005), this approximately corresponds to *c*_*LT M*_ = 0.27 (see Girard et al., 2023, for details). The ARM weights were initialized at 0; the LTM weights at 0.5.

An alternative energy model, termed *M*_0_, assumes that the energy is used whenever LTM plasticity occurs, but it is independent of the amount of synaptic strength change,

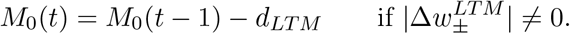

Simulation of the single exposure experiments, yields a calibration *d*_*LT M*_ = 0.1. Mathematically, the energy models corresponds to the *L*_1_ and *L*_0_ norms of the weight updates (van Rossum, 2023). The subscript distinguishes between the two variants for the energy consumed by LTM. To summarize

*M*_1_**-energy:** Energy equals the total amount of LTM synaptic weight change, e.g. number of receptors inserted and removed.

*M*_0_**-energy:** Energy equals the total number of LTM events.

We are not aware of experiments that decide between these energy models; future experiments hopefully will. Note that interactions between ARM and LTM pathways as well as interactions across time, that could in principle increase or reduce energy requirements, are also ignored.

### Stimulus protocol

In the simulation an odor is presented each day, which when approached, leads to a hazard of killing the fly and hence is to be avoided. In detail, on every day: the fly chooses stochastically to approach or avoid the stimulus (Eq. 4); the reward and reward expectation are updated (Eq. 7); the synapses are updated (Eq.5); the energy reserve is updated; the hazards are calculated; and finally the expected reward and ARM weights are decayed. The protocol is given to 10000 flies and repeated 50 days at the end of which all flies will have died.

The simulation contains in principle two stochastic elements: first, the decision to avoid the stimulus stochastic (Eq.4) and, second, the hazard is a probability to be evaluated every day for every fly, Fig.1. As a technicality, by calculating the population average expected lifetime from the hazards (Eq. 1), we remove this second source of variability in the simulation and reduce variability that would otherwise require larger simulated populations. Code for the paper can be found at github.com/vanrossumlab/neuroeconomicRL.

## Results

### ARM versus LTM learning

We first illustrate the model by assuming that flies exclusively use either the ARM or the LTM pathway. We simulate a population of flies that is subject to the following experiment: Each day an odor is presented, which when approached, leads to a hazard of killing the fly and hence is to be avoided. In addition there is a hazard to die from starvation, Eq. 2. We initially assume that apart from the energy required for LTM learning, there is no change in the energy reserve in the flies.

We track the evolution of the hazard, synaptic weights, energy reserve, and performance as measured by avoidance of the stimulus, Fig. 3a. With ARM-pathway learning (left panels), performance improves but does not exceed 70%, as the flies forget between the exposures. As a result the stimulus hazard remains substantial, however, the energy reserve stays high and starvation hazard low. The synaptic weights (and as a result behavior) oscillate slightly before settling down due to the updates in the expected reward.

**Figure 3.**
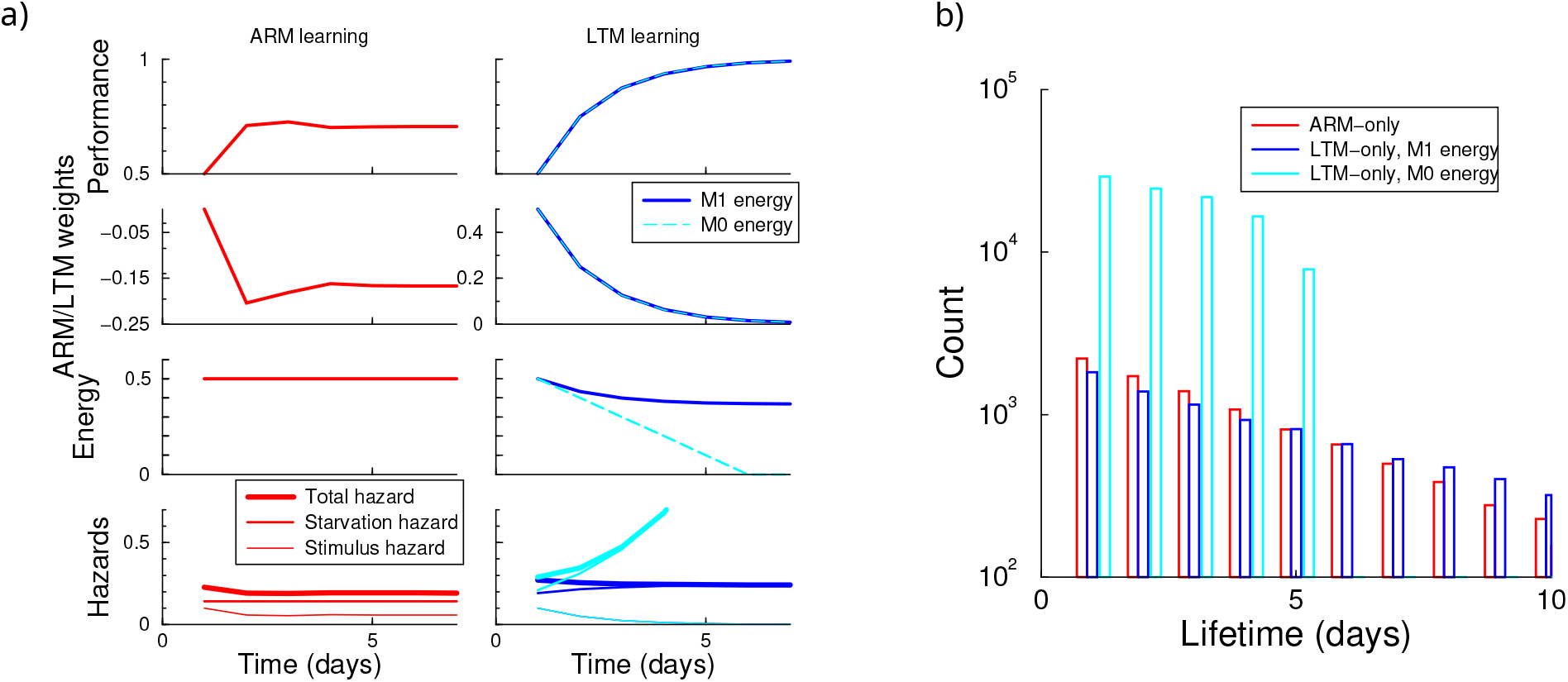
ARM vs LTM learning during the simulated aversive conditioning protocol. a) Evolution of performance, weights, energy reserve, and hazard. Left panel: ARM only learning. Right: LTM learning under the *M*_0_ and *M*_1_ energy model. b) Life-time histogram of 10000 flies under either pathway. Because the total hazard variations are relatively small, the distributions are close to exponential. However, for the *M*_0_ energy model, lifetimes are much shorter. (Parameters: stimulus hazard *h*_*s*_ = 0.2, initial energy reserve 0.5).

In contrast, in LTM learning performance becomes close to perfect after some 4 days, always avoiding the stimulus hazard, Fig. 3a right panels. The performance increases only slowly, because on the first trial only half the flies will randomly approach the stimulus and will learn, and so on.

While the hazard will be avoided, the expenditure of energy needed for LTM learning increases the starvation hazard. This effect is mild if the energy consumed by LTM is proportional to the size of the weight update (*M*_1_ energy). In this case the difference between the reward and its expectation and hence the amount of weight change diminishes as learning progresses. Only the first few learning events are costly (blue curves). However, when the cost is independent of the amount of weight update (*M*_0_, cyan curves), the energy is quickly depleted and the starvation hazard rises rapidly.

In this example ARM learning yields the longest lifetime of 5.6 days, LTM learning yields 4.4 days using the *M*_1_ energy model; their lifetime distributions are close to exponential. Using the *M*_0_ energy model the lifetime is only 2.4 days, Fig. 3b.

These simulations raise the question which memory pathway generally yields to longest life time for a given hazard and initial energy reserve. We varied the initial energy reserve of the flies, and determine the lifetime with ARM and LTM learning and in the absence of learning, Fig. 4a. Because in the model ARM learning comes at no cost, it is always better to learn with ARM than not learning at all. Under the *M*_0_ energy model, LTM learning never extends lifetimes (the cyan curve lies under all others). Under the *M*_1_ energy model (blue curve), there is a transition point. When initial energy is low, avoiding starvation is more important than avoiding the hazard, hence ARM yields longer lifetimes than LTM. In contrast, with a large energy reserve, the investment in avoiding the hazard is worthwhile and LTM yields a longer lifetime.

**Figure 4.**
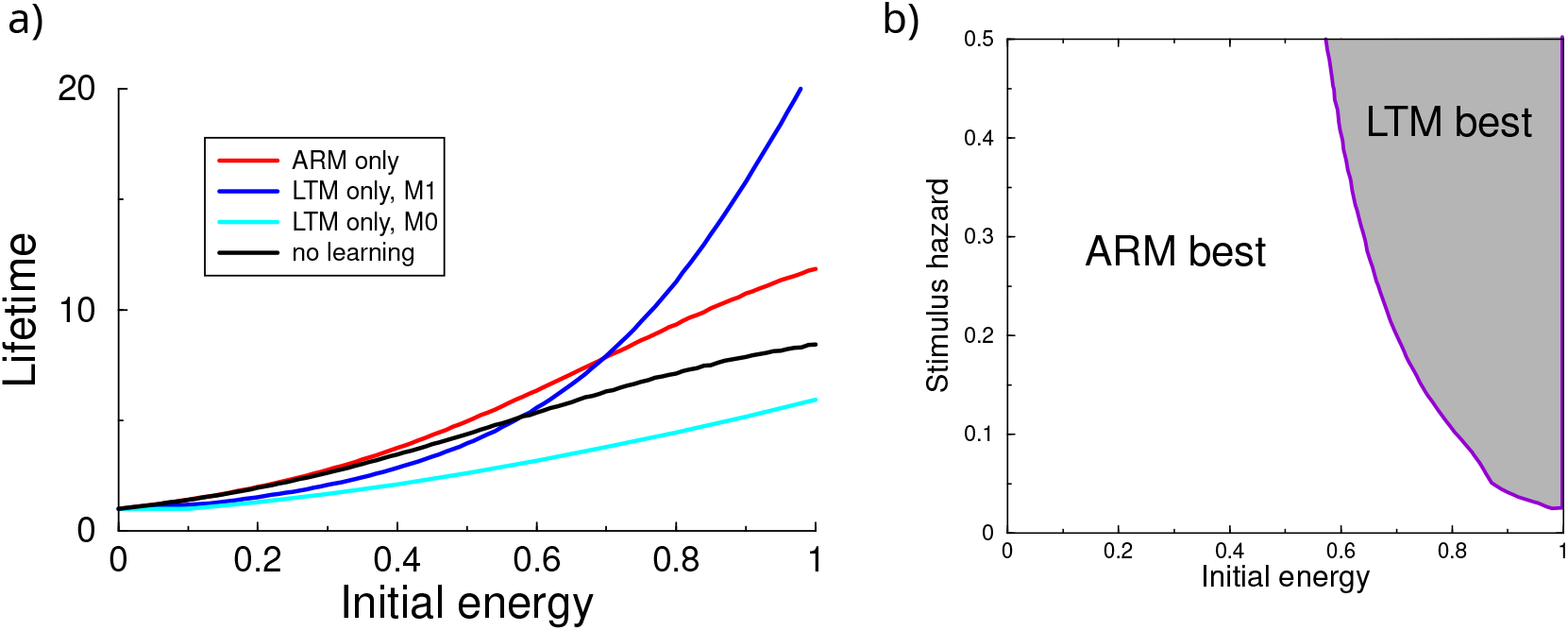
a) Life time effects of learning as a function of the energy reserve at day 0. ARM learning (red curve) is always better than no learning (black line). LTM (blue curve) is only beneficial when the energy reserve is high and the energy use is proportional to update size. (Stimulus hazard *h*_*s*_ = 0.1). b) Both the initial energy and hazard level influence whether LTM learning increases life-time over ARM learning. When the hazard is high, is better to invoke LTM at lower energy reserves (*M*_1_ energy model).

The point at which LTM is better, depends on the stimulus strength. The higher the hazard, the lower the transition point, Fig. 4b. In the Appendix we derive an equation that gives insight in the break-even point, however, a full analytical treatment seems out of reach, because learning does not only affect the next decision, but all future (discounted) decisions. In the remainder we therefor rely on simulations.

### Threshold Models

In the above simulations the memory pathway was set once and for all at the start of the simulation. While this is useful to gain understanding, it makes more sense to choose the pathway depending on the *current* energy reserve *M* (*t*). We assume that the expensive LTM pathway was used whenever the energy reserve exceeded a threshold, otherwise the ARM pathway was updated. To show the benefit of this algorithm we consider a population of flies with different initial energies, drawn uniformly between 0 and the maximum and measured the average lifetime as the threshold was varied, Fig. 5. Note that, as expected, when the threshold is 0 (1), the lifetime equals that of LTM-only (ARM-only). When the threshold parameter is tuned (x-axis), the lifetime can exceed that of exclusively using either LTM (blue) or ARM (red). The peak of the curve shifts left as the stimulus hazard increases. That is, the larger the stimulus hazard, the lower the threshold. Under the *M*_0_ energy model, adaptive learning is only beneficial for large stimulus hazards, Fig. 5 bottom row.

**Figure 5.**
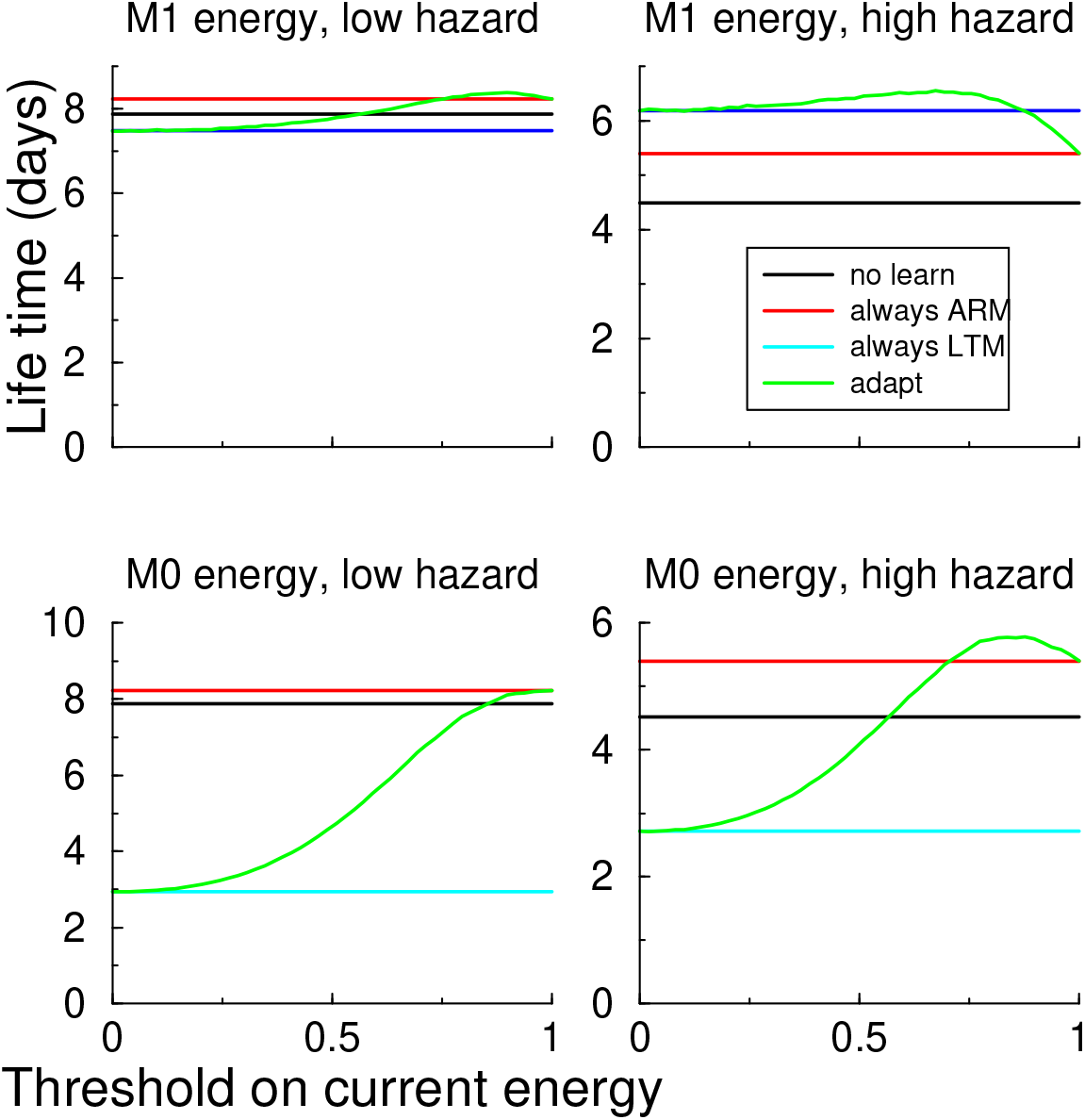
Adaptive switching between ARM and LTM can improve lifetime. Population life time vs threshold for two hazard levels. LTM was employed whenever the energy exceeded a certain threshold (x-axis). Left: low hazard (0.05) and high hazard (0.2). Using the *M*_1_ energy, the adaptive model (green) increases average population life time compare d to either LTM or ARM exclusively. For the *M*_0_ energy model this only happens at large hazards. The optimal threshold that gives the longest lifetime depends on stimulus hazard.

### General threshold models

In the above adaptive switching model, LTM will be employed when the energy reserve is sufficient, even if the reward prediction error and hence weight changes are small. This means that energy might be spend for only a small change in avoidance behavior. Therefore, we made the threshold both dependent on the current energy reserve *M* (*t*), and the difference between expected and actual reward,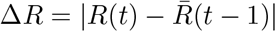. We parameterized the switch so that the LTM pathway was employed whenever

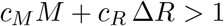

The parameters *c*_*M*_ and *c*_*R*_ define a line the *M*, Δ*R*-plane. When *c*_*R*_ is set to zero, we retrieve the energy threshold model: *M* has to be larger than 1*/c*_*M*_ for LTM to occur, Fig 6a. Likewise, when *c*_*M*_ = 0 the decision solely depends on Δ*R*. Generally, when *c*_*M*_ *>* 0, a large reward prediction error Δ*R* will lower the threshold for LTM memory.

**Figure 6.**
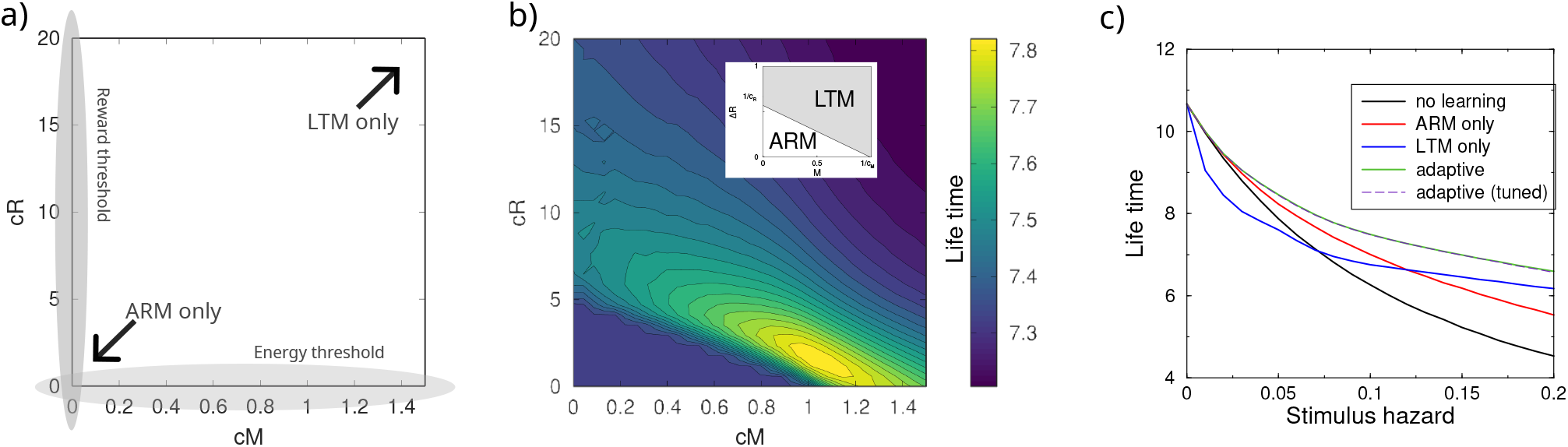
Adaptive plasticity model with dependence on both the current energy reserve and the reward prediction error and parameters *c*_*M*_ and *c*_*R*_. LTM was only used when *c*_*M*_ *M* + *c*_*R*_ |Δ*R*| ≥1. b) Life-time as a function of the threshold parameters. Life time was averaged across stimulus hazards. Inset show the corresponding optimal threshold model. c.) Life-time as function of the stimulus hazard. The adaptive plasticity yields the longest life time. (M1 energy model)

**Figure 7.**
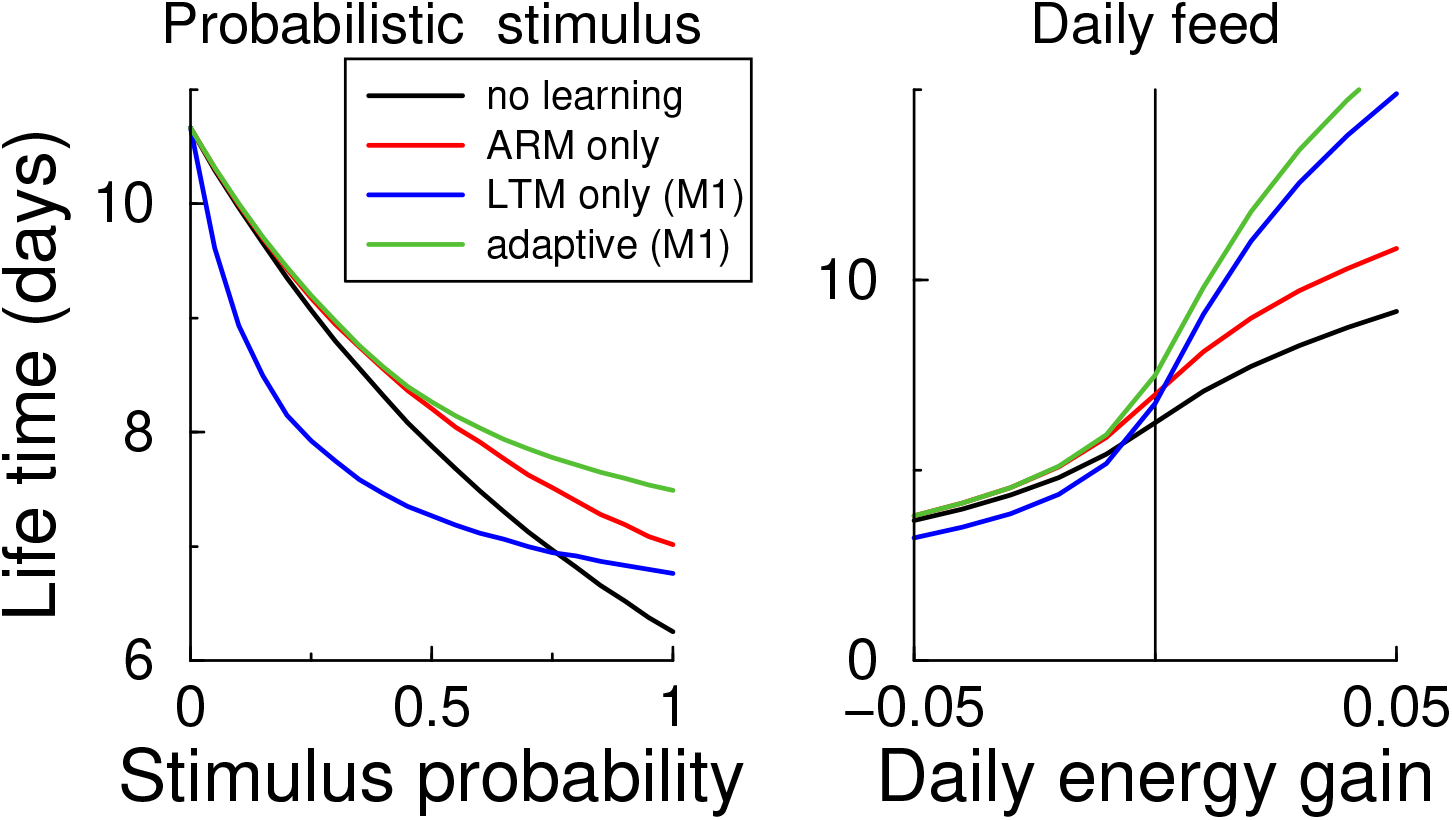
The life times for a probabilistic stimulus (left) and when there is additional intake or loss (right). The adaptive algorithm robustly outperforms fixed strategies (Stimulus hazard 0.1; *M*_1_ energy model)

We varied the stimulus hazard and optimized the *c*_*M*_ and *c*_*R*_ parameters of the threshold. The life-time is maximal around *c*_*M*_ = 1.01 and *c*_*R*_ = 1.76, Fig. 6b. If a threshold on just the energy were optimal, the optimal threshold would be lying on the *c*_*R*_ = 0 axis. And similarly, when just a threshold on Δ*R* would suffice, the optimal solution would lie on on the *c*_*M*_ = 0 axis. As the optimum lies away from both axes, a joint threshold yields the longest lifetimes.

The lifetime using optimized parameters exceeds that of exclusively using the ARM or LTM pathway across stimulus hazards, Fig. 6 right. The adaptive threshold model picks the ‘best of both worlds’.

Ideally, the optimal threshold should be such that a change in stimulus hazard should not require a re-tuning of the threshold parameters. We calculated the life-time for the parameters that were best on average, and compared it to the life-time optimized for each value of stimulus hazard. The lifetimes using the fixed parameters were practically indistinguishable from the individually tuned threshold parameters. Hence the threshold model is robust against changes of the stimulus parameter..

We also tried a variant in which either energy reserve *or* reward error where above a threshold, as well as a model where both energy reserve *and* reward error needed to exceed a threshold; these both did not perform as well as the above model (not shown).

For the *M*_0_ energy model the results are very comparable (Supplementary fig 9). As above, under the *M*_0_ energy model the lifetime is severely shortened when always using the LTM pathway, because every LTM plasticity is expensive even if the weight changes are small. But again, adaptively switching to LTM under the right circumstances again improves lifetime, Fig.9a. The optimal *c*_*M*_ parameter is somewhat smaller (*c*_*M*_ = 0.97, *c*_*R*_ = 2.35), that is, the energy needs to be larger to switch to LTM than for the *M*_1_ energy model, Fig.9 b.

We repeated this analysis for two other parameters of the stimulation protocol. First, we fixed the stimulus hazard (0.1), but we assumed that approaching the stimulus only sometimes lead to exposure to the hazard. The hazard probability was varied between 0 and 1, and determined on each trial independently whether the hazard was encountered or not. As expected, at the zero stimulus probability, the lifetime was maximal and independent of any learning. Again the adaptive threshold robustly improved lifetime, Fig.7 left.

Next, we modified that model so that in addition to the energy consumption by LTM plasticity, there was a fixed daily energy intake/consumption. The lifetime has a sigmoidal shape as a function of this amount, Fig.7 right. When there is a high consumption (left part of graph), the fly heads for perishing anyway, and investment in LTM learning only hastens that. But when there is a daily net intake, the investment in LTM memory helps to escape the hazard, while future starvation is unlikely. The lifetimes using LTM memory now exceed those from ARM learning. Again the adaptive algorithm improves the lifetime, outperforming either ARM or LTM exclusive learning. The results for both daily energy intake/consumption and the probabilistic stimulus also hold when considering the *M*_0_ energy (not shown).

### Appetitive conditioning

While we designed the model for avoidance conditioning, the same circuit is thought to underlie appetitive conditioning. To model this we assumed that approaching the reward increased the energy reserve with an amount 0.05. The only hazard that the fly encountered was from starvation. We first examined the effect of learning on life time as a function of initial energy for the various learning protocols. In contrast to aversive conditioning, the learning makes only little difference on the life time, Fig. 8left, cf. Fig.4. ARM learning performs very well throughout. The reason is that in contrast to the aversive protocol, the ARM memory is daily refreshed by approaching the stimulus and boosted in appetitive conditioning. Moreover, as the maximum energy was capped at 1, gaining extra reward did not carry a large benefit, whereas in aversive conditioning avoiding the hazard is always important.

**Figure 8.**
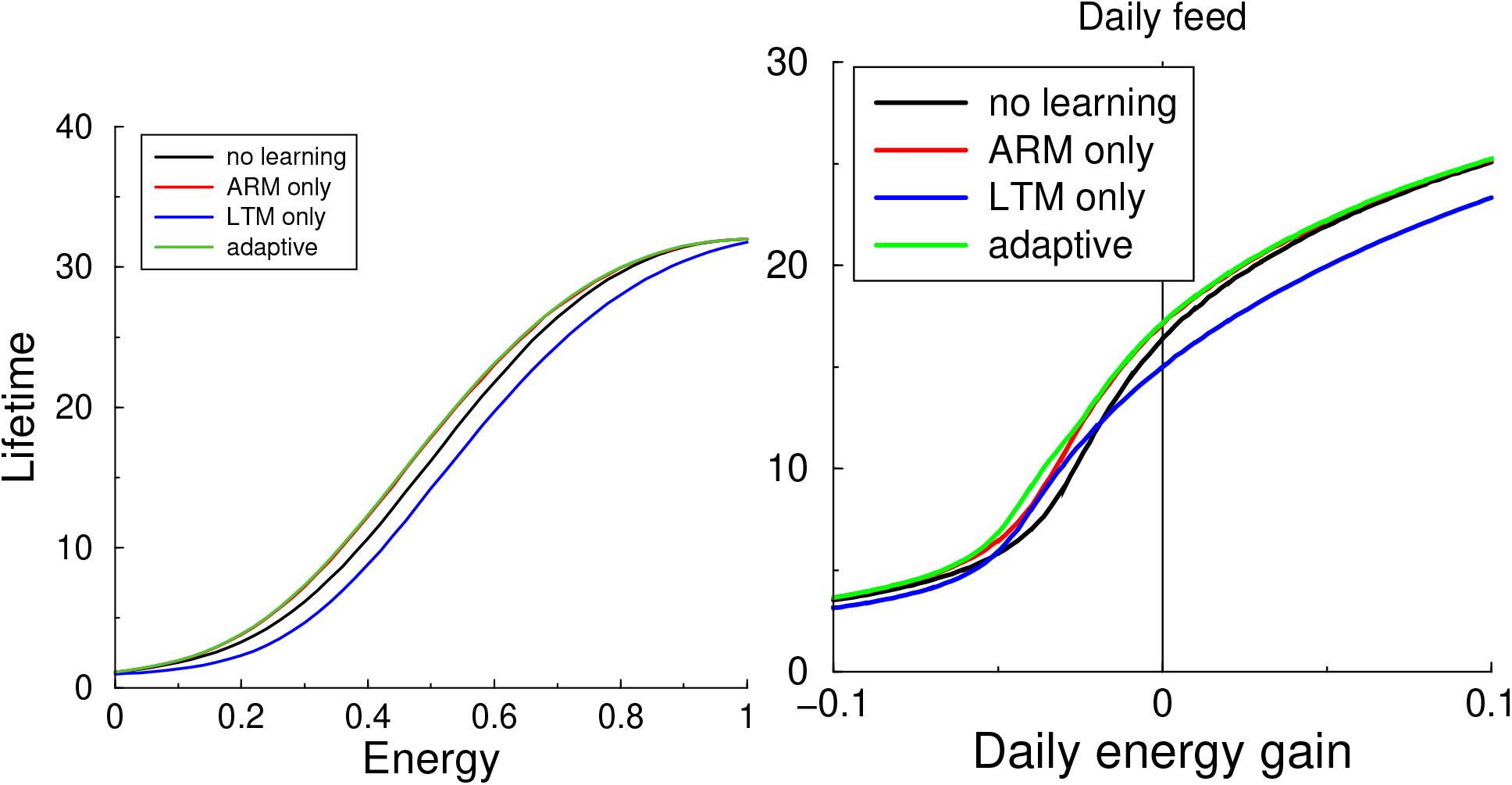
Appetitive conditioning. a) The lifetime under appetitive conditioning as a function of the initial energy reserve. LTM performs worse, while ARM performs close to optimal. b) Life-time as a function of daily intake. While adaptive learning is still best, differences are again small.

**Figure 9.**
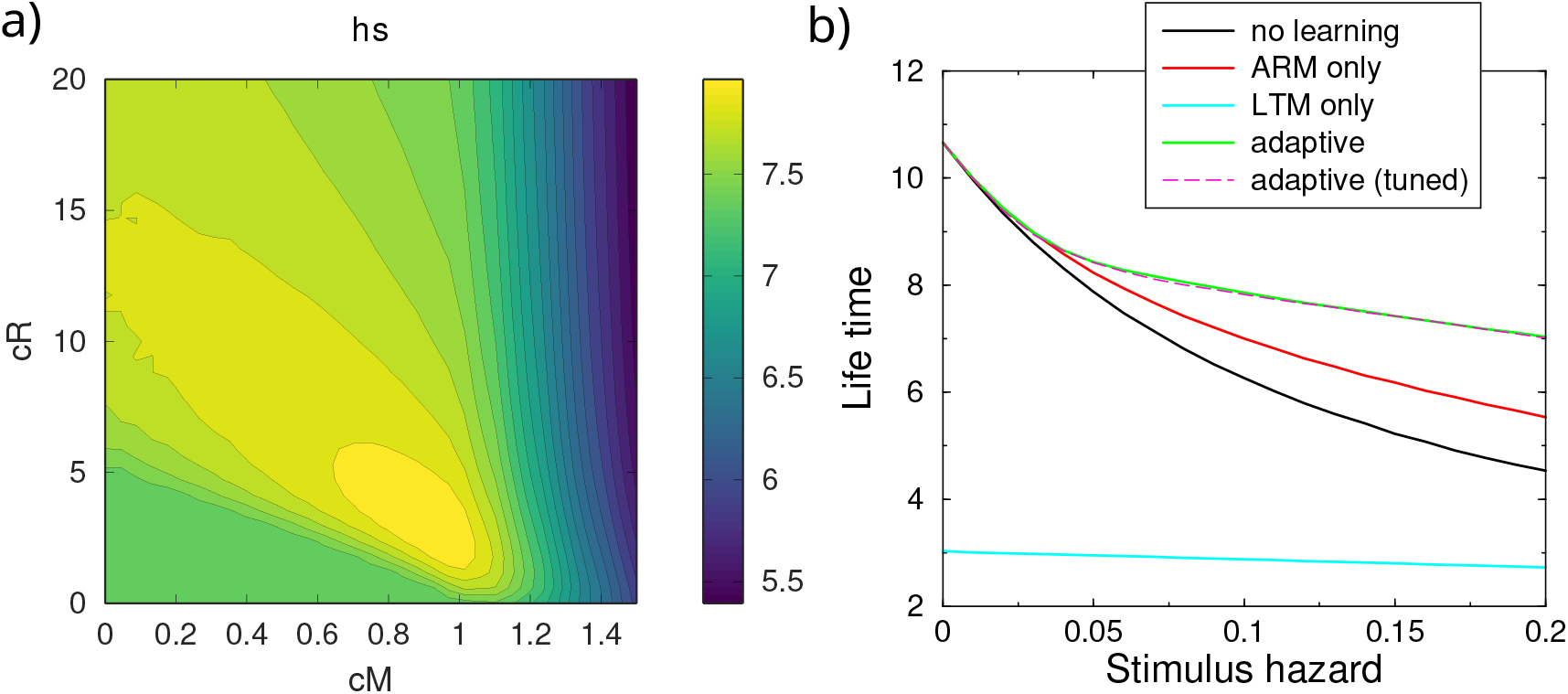
Supplementary figure. As Figure 6 but using the *M*_0_ energy model. a) Life-time as a function of the threshold parameters. Average across stimulus hazards b) Life-time as function of the stimulus hazard. As for the *M*_1_ energy model, the adaptive plasticity yields the longest life time.

Next, we included a daily net energy increase or decrease. As in the above aversive conditioning, the mean life time depends strongly on this daily amount, Fig. 8 right. Learning however only slightly improves lifetime. Furthermore, for the used parameters LTM learning was never beneficial over ARM learning. Only in a narrow region LTM learning outperforms no-learning.

It is known that flies also switch between LTM and ARM pathways in appetitive conditioning. However, in contrast to aversive conditioning, the LTM pathway is only activated when the animals are starved prior to conditioning (Trannoy et al., 2011). Unlike the experimental findings, there would seem no reason why LTM would more beneficial at low energy than at high energy reserves, Fig. 8left. When the animal has enough reserve there is no reason not to use LTM. However, it might that the LTM requires other resources that are scarce, or that LTM learning carries other detrimental consequences.

## Discussion

Inspired by experimental findings that LTM memory formation is metabolically costly, and that flies stop aversive LTM learning under starvation, we have explored how such adaptive learning can increase evolutionary fitness and how the switch between LTM and ARM should be set. Using the hazard framework, a switch to LTM memory when the energy reserve is high and the reward prediction error is high, improves population life-time.

Necessarily, simulations need to assume a certain hazard exposure protocol. The optimal parameters that set the switch point will be dependent on this. But some generalizations are immediately obvious. For instance, when the stimulus interval is increased, the ARM memory will decay more between events, and ARM becomes less effective. As a result the fly should switch to LTM sooner. As another extreme example, if the stimulus were only presented once, learning would be useless and should be turned off. The biological parameters have presumably been optimized for performance across the ensemble of naturally encountered environments and hence the parameters values found here are not expected to be exactly those found in experiments. Future studies could aim to close this gap and study more realistic and richer environments, including those with temporal correlations. The adaptive algorithm might be adjusted to include stimulus repetition and spacing effects (Anderson and Schooler, 1991).

We have relied on mean population lifetime as fitness measure, however true fitness is the ability to pass genetic material to offspring. A more involved model could use a fitness measure that reflects that. For instance, for a population it might be better to have a wide spread in the life-time distribution, so that some individuals would survive periods of famine.

While the current work focused on Drosophila anatomy and physiology, there are indications that similar principles might be at work in mammals. In contrast to the fruit-fly’s ARM and LTM pathways, transient and persistent forms of mammalian long term potentiation (LTP) appear to be expressed at the same synapse. However, also in mammals there is physiological evidence for down-regulation of persistent LTP under energy scarcity via the AMPK pathway (Potter et al., 2010), and there is behavioral evidence for a correlation between blood glucose level and memory formation (Gold, 1986; Smith et al., 2011). Likewise, the presence of a dopamine reward signal, typically interpreted as signaling the reward prediction error, lowers the threshold for late-phase LTP (O’Carroll et al., 2006; Bethus et al., 2010; Lisman et al., 2011). Finally, reinforcement learning has many engineering and software applications. The results found here could potentially enhance the performance of RL algorithms, especially in resource-limited settings or tasks requiring multi-objective optimization. The energy requirements in these applications could be associated to computing the weight updates, but also for computer hardware, memory storage is energetically expensive.

## Acknowledgments

JJ was supported by a Nottingham Vice Chancellor’s International Scholarship, and a Turing Institute Enrichment scheme R00700. Discussions with Aaron Pache, Silviu Ungureanu, Long Tran-Thanh, Andrew Lin, and Walter Senn are gratefully acknowledged.

## 1 APPENDIX Micro economic Trade-off for learning

The optimal memory strategy maximizes the lifespan. The animal has to decide whether to invest energy in long term memory of the CS-US associate. We derive an expression for the change in lifetime given a small weight update, which in turn leads to a small change in the hazards *δh*(*t*). Under this assumption the change in expected lifetime between LTM and no learning (NL) can be expanded as

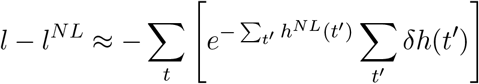

Given ARM learning with a small weight change Δ*w*, the temporary reduction in stimulus hazard is

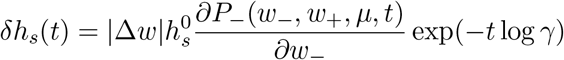

For LTM learning, the expression is similar but the decay term is absent. LTM learning at the same time increases starvation hazard as

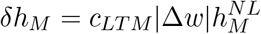

The difference in expected lifetime between ARM and LTM learning is in first order of |Δ*w*|,

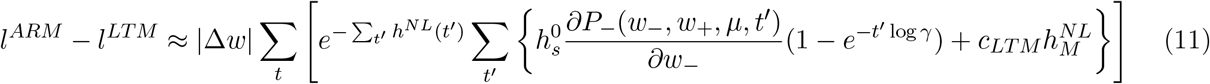

Where it should be noted that because learning decreases the probability of encountering the stimulus (*∂P/∂w <* 0), the first term in the curly brackets is negative, while the second term is strictly positive. 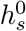 denotes the stimulus hazard if it is approached.

When the lifetime difference is larger than zero, ARM learning should be chosen over LTM learning. While complex, the expression gives insight in when ARM memory is preferable to LTM. It happens when: 1) The stimulus hazard 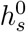 is small, 2) when the impact of the learning on the choice probability *∂P/∂w* is small, e.g. late in the learning process, 3) the ARM decay *γ* is slow, and 4) the energy cost of LTM, *c*_*LT M*_ is high. Finally, the first r.h.s term attenuates the benefit of long lasting memory, so that ARM is generally preferable when the expected lifetime is short.

Nevertheless, it would appear challenging for a fly to estimate the expected life time based on this expression to decide whether to use ARM or LTM memory, so instead we a looking for approximate heuristic algorithms that only rely on observables directly accessible by the organism and are close to optimal under various conditions.

