## Supplementary material for "Reinforcement learning when your life depends on it: a neuro-economic theory of learning": Supp Fig

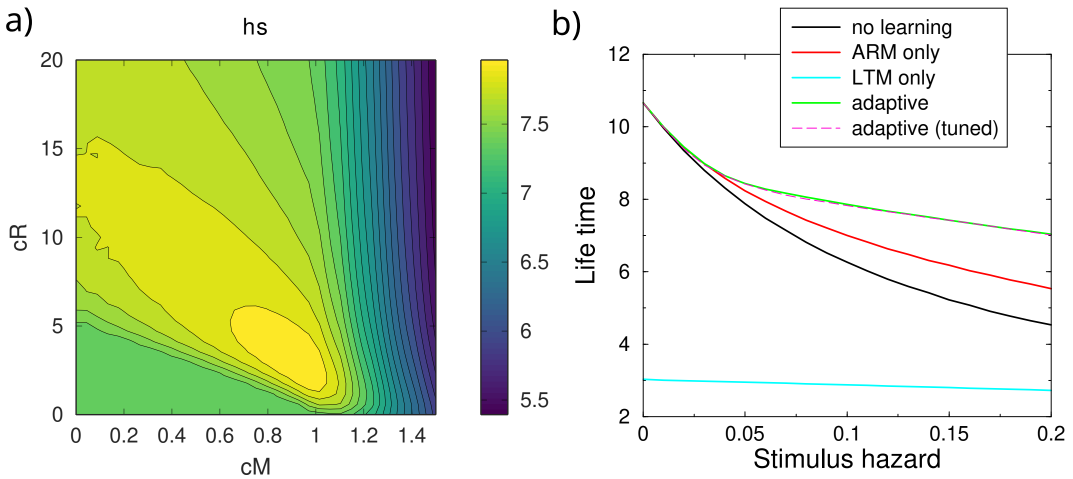


Figure 9: Supplementary figure. As Figure 6 but using the $M_0$ energy model. a) Life-time as a function of the threshold parameters. Average across stimulus hazards b) Life-time as function of the stimulus hazard. As for the $M_1$ energy model, the adaptive plasticity yields the longest life time.
